# Higher anhedonia during withdrawal from initial opioid exposure is protective against subsequent opioid self-administration in rats

**DOI:** 10.1101/839993

**Authors:** Yayi Swain, Peter Muelken, Annika Skansberg, Danielle Lanzdorf, Zachary Haave, Mark G. LeSage, Jonathan C. Gewirtz, Andrew C. Harris

## Abstract

Understanding factors contributing to individual differences in vulnerability to opioid addiction is essential for developing more effective preventions and treatments, yet few reliable behavioral predictors of subsequent opioid self-administration have been identified in rodents. Sensitivity to the acute effects of initial drug exposure predicts later addiction vulnerability in both humans and animals, but the relationship of sensitivity to *withdrawal* from initial drug exposure and later drug use vulnerability is unclear. The goal of the current study was to evaluate whether the degree of anhedonia experienced during withdrawal from early opioid exposure predicts subsequent vulnerability to opioid addiction. Rats were first tested for withdrawal sensitivity following acute injections of morphine (i.e., “acute dependence”), measured as elevations in intracranial self-stimulation (ICSS) thresholds (anhedonia-like behavior) during naloxone-precipitated and spontaneous withdrawal. Rats were then tested for addiction vulnerability using various measures of i.v. morphine self-administration (MSA) including acquisition, demand, extinction, and reinstatement induced by morphine, stress, and/or drug-associated cues. Greater naloxone-precipitated withdrawal across repeated morphine injections and greater peak spontaneous withdrawal severity following a single morphine injection were associated with lower addiction vulnerability on multiple MSA measures. Withdrawal-induced anhedonia predicted a wider range of MSA measures than did any individual measure of MSA itself. These data suggest that high anhedonia during withdrawal from initial opioid exposure is protective against subsequent opioid addiction vulnerability in rodents, thereby establishing one of the first behavioral measures to predict individual differences in opioid SA. This model promises to be useful for furthering our understanding of behavioral and neurobiological mechanisms underlying vulnerability to opioid addiction.

## 1. Introduction

Opioid addiction poses a substantial burden on public health [1, 2]. Identifying behavioral and neurobiological factors contributing to the marked individual differences in opioid addiction vulnerability is essential for developing more effective preventions and treatments [3, 4]. However, few behavioral measures have been identified in preclinical models that reliably predict individual differences in i.v. opioid self-administration (SA). SA is often considered a “gold standard” for modeling addiction in animals because it involves volitional drug consumption as occurs in humans.

Sensitivity to the initial acute effects of drugs (e.g., euphoria, aversion) is a key predictor of addiction vulnerability in humans [5–7]. Furthermore, several preclinical studies indicate that sensitivity to the initial effects of acute drug injections (e.g., locomotor activity or depression, antinociception) predict voluntary drug intake in an i.v. SA model [8–10]. Acute drug injections also produce negative affective (emotional) states (e.g., anhedonia, or diminished reward sensitivity) during withdrawal. These withdrawal effects are induced even after a single drug exposure in both humans and animals (“acute dependence”; [11, 12, 13]), and often become more severe with repeated drug exposures [14–16]. Some authors have proposed that greater sensitivity to the negative affective consequences of withdrawal may be protective against addiction [17–21], and that anhedonia may reduce the motivation for reward-seeking [22]. Consistent with these predictions, we found that saccharin-preferring rats, which exhibit greater SA of opioids and other drugs [23], exhibit lower anhedonia during morphine withdrawal as measured by increases in intracranial self-stimulation (ICSS) thresholds (i.e., withdrawal-induced anhedonia, WIA) [19]. Nevertheless, the relationship between sensitivity to withdrawal from acute opioid exposure and opioid SA has not been directly tested within the same animals or in outbred rats, which can differ from selectively bred rats in terms of determinants of addiction vulnerability [24, 25].

The goal of this study was to evaluate the ability of WIA to predict individual differences in subsequent i.v. morphine SA (MSA) in outbred rats. Several different measures of MSA (e.g., acquisition, demand, reinstatement) were used because they model distinct aspects of addiction and can be differentially associated with other behavioral predictors of drug SA [26, 27]. It was hypothesized that greater WIA severity would be associated with lower MSA vulnerability.

## 2. Materials and Methods

### Overview of experimental protocol

Male adult Sprague-Dawley rats (see Supplementary Material for additional information) were tested under the experimental protocol shown in Figure 1. Rats were first tested for WIA during naloxone-precipitated and spontaneous withdrawal from acute morphine injections (Phase 1). Rats were then tested for opioid addiction vulnerability using several measures of i.v. MSA including acquisition, elasticity of demand, and reinstatement (Phase 2). Finally, WIA was again tested after the completion of the MSA protocol to provide a *preliminary* characterization of the relationship between MSA and withdrawal sensitivity during a more advanced stage of dependence (“late-stage dependence”, Phase 3). All procedures were approved by The Institutional Animal Care and Use Committee (IACUC) of the Hennepin Health Research Institute in accordance with the 2011 NIH Guide for the Care and Use of Laboratory Animals and the 2003 National Research Council Guidelines for the Care and Use of Mammals in Neuroscience and Behavioral Research.

**Figure 1.**
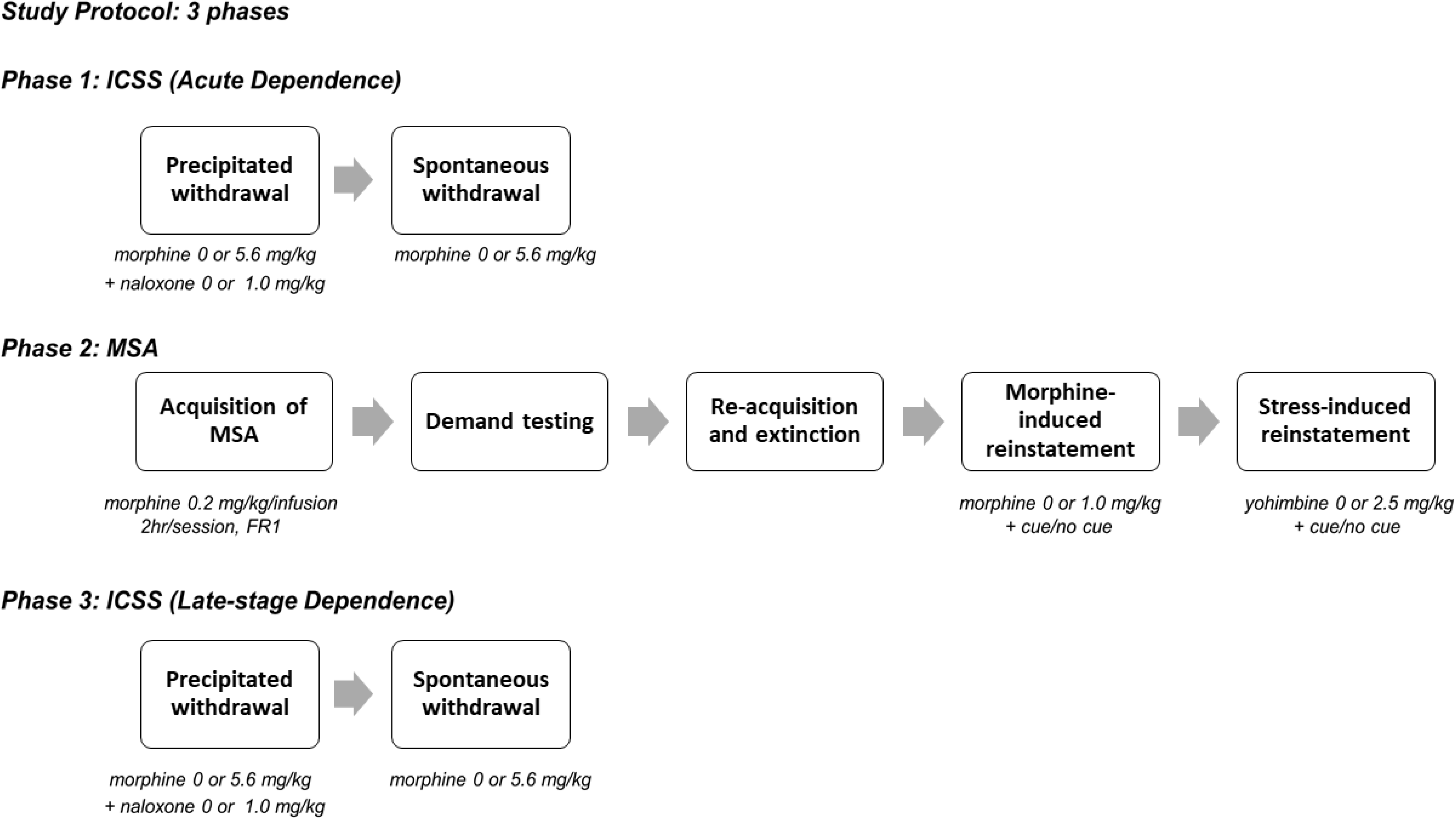
Overview of experimental protocol (see text for more details).

### Acute dependence

Rats (N = 61) were prepared and trained on a discrete-trial ICSS procedure (see Supplemental Material) in daily sessions conducted Mon-Fri until ICSS thresholds were stable (<10% variability over 5 days) and habituated to saline injections as described previously (see [28]). On the first test day, rats were injected with morphine sulfate (MOR, 0 or 5.6 mg/kg, s.c., expressed as the salt). One hour and fifty minutes later, rats were injected with the opioid antagonist naloxone (NX, 0 or 1.0 mg/kg, s.c.) and tested for ICSS 10 minutes later. These morphine and naloxone doses and this pretreatment interval produce significant negative affective morphine withdrawal signs, including WIA [12, 15, 19]. Immediately after ICSS testing, somatic withdrawal signs were assessed as a secondary withdrawal measure (see Supplementary Material). There was a total of 4 groups in a 2 (MOR dose) x 2 (NX dose) factorial design. The MOR + NX group (n = 29) was larger than the MOR + SAL, SAL + NX, and SAL + SAL groups (n = 10-11/group) in order to have adequate power for correlation analysis (see below). These procedures were repeated each day for 5 consecutive days. Animals were then tested for ICSS under drug-free conditions for at least one week and until ICSS thresholds were stable (same stability criteria as above).

To test spontaneous withdrawal, rats received a single injection of 5.6 mg/kg MOR (MOR+NX and MOR+SAL groups) or 0 mg/kg MOR (SAL+NX and SAL+SAL groups) and ICSS was tested 2, 6, 26, 30, 50, 64, 74, 98, and 170 hours later. The purpose of the 2 hour time point was to detect any reinforcement-facilitating (ICSS threshold-lowering) effects of morphine itself (see [29]). The selection of subsequent time points was based on the time course of spontaneous withdrawal from acute morphine exposure determined using ICSS and other measures [11, 30, 31]. Somatic withdrawal signs were recorded after ICSS testing at only the 26 hour time point. ICSS testing was then suspended.

### Locomotor activity

Within 48 hours after the final ICSS test, locomotor activity in a novel environment (i.e., “sensation-seeking” [32–34]) was tested for 2 hours as a secondary predictor of MSA (see [35] for detailed description of the apparatus).

### MSA

Approximately 24-48 hours after completion of the locomotor activity test, animals from all groups were implanted with i.v. catheters using our standard procedures [34]. Rats from the control groups (SAL + SAL, etc.) were included in this phase to determine whether a history of withdrawal testing impacted subsequent MSA in the MOR + NX group. Following a 7-10 day recovery period, all rats were allowed to acquire i.v. morphine SA (0.2 mg/kg/inf) during daily 2 hr sessions conducted Mon-Fri using our standard apparatus and procedures (see Supplementary Material). Rats were tested under a fixed ratio (FR) 1 schedule for at least 10 sessions and until acquisition criteria were met (≥5 infusions per session, ≤20% coefficient of variation, and ≥ 2:1 response ratio on the active lever to inactive lever) across 3 sessions. To test elasticity of demand (reinforcing efficacy), the FR requirement was increased every 3-4 sessions as follows: FR 2, 3, 6, 12, 24, and doubled thereafter until infusion rates during the last 2 sessions at a given FR were reduced by 90% compared to baseline (FR 1). Morphine consumption under this protocol is well described by the current exponential demand function [35]. Data from Mondays were excluded from data analysis due to spontaneous recovery of responding after the weekend, which was often observed at high unit prices. Therefore, if one of the three sessions at a given FR occurred on a Monday, rats were tested in an additional session at that FR.

After completion of demand testing, rats were allowed to reacquire MSA under an FR1 schedule for at least 5 sessions and until MSA was stable (same stability criteria as above). Extinction conditions were subsequently introduced in which the morphine dose was replaced with saline and the drug-associated cue light was no longer presented upon infusion. Extinction was tested for at least 10 sessions and until animals exhibited a 75% reduction in active lever pressing for 2 consecutive sessions.

To test morphine- and cue-induced reinstatement, rats were injected s.c. on separate days with either saline or morphine (1.0 mg/kg) 10 min prior to the SA session. This morphine dose and pretreatment interval reliably reinstated drug-seeking in our lab (data not shown) and others [36]. Responses on the active lever resulted in a saline infusion and either presentation of the drug-associated cue light (“cue” condition) or no programmed consequences (“no cue” condition). Reinstatement testing was therefore conducted using a 2 (morphine dose) x 2 (cue condition) design, resulting in a total of 4 reinstatement tests. These tests were conducted on Tuesdays and Fridays, provided that active lever pressing returned to extinction levels during the preceding session, and the order of reinstatement conditions was counterbalanced. Following completion of morphine- and cue-induced induced reinstatement testing, rats were tested under extinction conditions for at least 5 sessions and until extinction criteria were again met. To test stress- and cue-induced reinstatement, the above procedure was repeated except that rats were injected i.p. with either deionized water or the stress-inducing α2-adrenergic antagonist yohimbine (2.5 mg/kg) 30 min prior to each SA test. This dose of yohimbine and pretreatment interval reliably reinstate extinguished SA [37].

### Late-stage dependence

Following completion of all MSA procedures, rats were again tested for ICSS until thresholds were stable. Rats (N = 26) were subsequently tested for precipitated (MOR + NX: n = 14, n = 4/group for other groups) and spontaneous (MOR + NX: n = 11, n = 2-4/group for other groups) morphine withdrawal as described above. The small group sizes for this exploratory phase reflect the considerable attrition rate by this stage of the protocol (see below).

### Statistical analysis

#### ICSS

ICSS thresholds (a measure of brain reinforcement function) and response latencies (a measure of non-specific motoric effects [38]) during naloxone-precipitated and spontaneous withdrawal were measured as percentage of baseline (average of the last 5 days prior to onset of withdrawal testing). To provide a composite measure of naloxone-precipitated withdrawal severity for each animal, standardized composite z-scores were determined based on average and peak ICSS thresholds during all 5 precipitated withdrawal tests, as well as degree of sensitization of WIA (the difference score in ICSS thresholds between test days 1 and 5). Pearson’s correlation was used to confirm that composite z-scores correlated significantly with all individual measures of WIA [see 39, 40]. Spontaneous withdrawal severity was measured as peak withdrawal severity during hours 6 – 98 after morphine injection. This measure accounts for the considerable individual differences in the time course of changes in ICSS thresholds during spontaneous withdrawal [see 41]. In the few cases where rats failed to respond for any ICSS current intensity, we arbitrarily assigned ICSS threshold and latency values based on those obtained in the animal achieving the highest ICSS threshold in that phase of the experiment [see 42, 43].

#### MSA

MSA acquisition was measured as the mean number of infusions per session during the first 10 days of acquisition. To determine opioid reinforcing efficacy during FR escalation, exponential demand curve analyses were conducted as described in detail elsewhere [35, 44]. Our primary demand measure, α, refers to the rate of change in consumption with increases in unit price (elasticity of demand), with higher α values indicating lower reinforcement efficacy. Zero values in consumption were replaced with 0.01 (1/10^th^ of our lowest non-zero consumption level) to provide better curve fits and more accurate parameter estimates of demand for individual rats [see 33, 45, 46]. α values were log-transformed due to non-normal distribution. Extinction was measured as mean number of infusions per session during the first 10 sessions of extinction. Degree of reinstatement was defined as the difference between active and inactive lever responses during each reinstatement test. To control for non-specific active lever pressing, reinstatement scores during the control condition (Vehicle + No cue) were subtracted from the reinstatement scores for the other conditions (Morphine + Cue, etc.) for correlational analyses.

All statistical analyses and graphing were performed in GraphPad Prism 7 or R 3.4.3, with significance level set at α = 0.05 for all tests. In general, data were analyzed using ANOVA followed by Holm-Sidak’s or Dunnet’s multiple comparison tests (see Results for more details). Relationships between ICSS and MSA measures were assessed using Pearson’s correlation.

## 3. Results

### Attrition

Several animals were lost to attrition during the course of the protocol due to loss of ICSS headcap, loss of stability of ICSS thresholds, failure to acquire MSA, loss of catheter patency, health issues, or other problem. Data for these animals are analyzed only for those phases they completed.

### Acute dependence

#### Precipitated withdrawal: ICSS

Baseline ICSS thresholds did not differ between groups (Table S1). A two-way ANOVA on ICSS thresholds during precipitated withdrawal revealed a significant main effect of group (F (3, 54) = 36.62, p < 0.0001), a marginally significant main effect of session (F (4, 216) = 2.34, p = 0.06), and a significant interaction between group and session (F (12, 216) = 3.181, p = 0.0003). Dunnett’s multiple comparisons indicated that ICSS thresholds were significantly elevated in the MOR+NX group compared to SAL+SAL controls during all 5 sessions (all q ≥ 3.162, all p ≤ 0.005). In contrast, thresholds in the MOR+SAL and SAL+NX groups did not differ from the SAL + SAL group during any session (Figure 2A). A repeated-measures ANOVA showed a significant effect of session in the MOR+NX rats (F (2.863, 77.30) = 18.40, p < 0.001), with Dunnett’s multiple comparisons indicating significantly higher ICSS thresholds during sessions 2-5 compared to session 1 (all q ≥ 4.68, all p ≤ 0.001). In contrast, there was no effect of session in any of the control groups (all p > 0.05).

**Figure 2.**
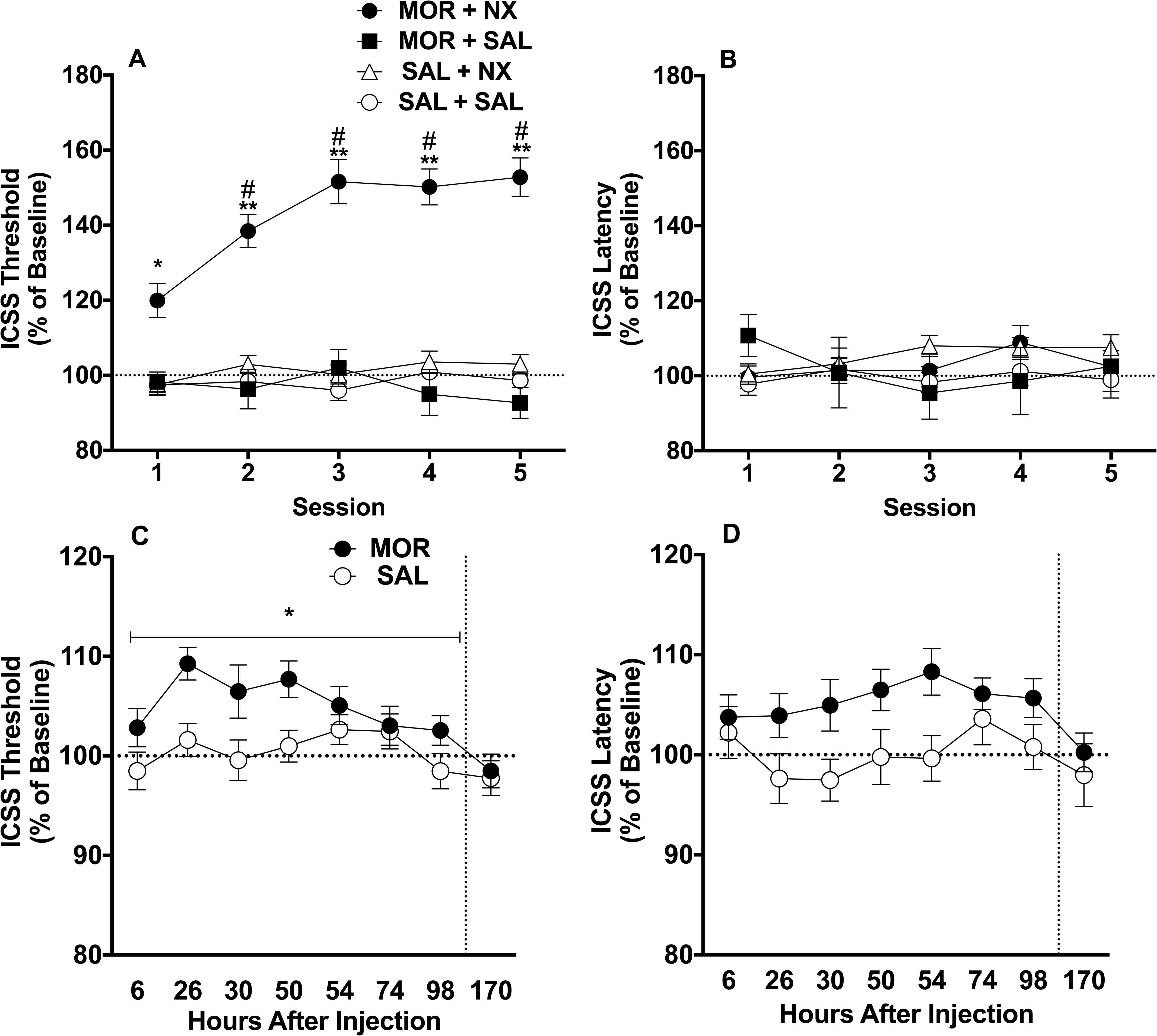
Mean (± SEM) ICSS thresholds (A) and response latencies (B) (expressed as percent of baseline) during naloxone-precipitated withdrawal (acute dependence). Mean (± SEM) ICSS thresholds (C) and latencies (D) as percent of baseline during spontaneous withdrawal (acute dependence). *,** Different from SAL+SAL group at that session or SAL condition during hours 6-98 (main effect), p < 0.05, 0.01. # Different from Session 1 in that group, p < 0.05.

No significant differences were observed in baseline ICSS response latencies between groups (Table S1). Latencies also did not differ between groups during precipitated withdrawal testing (Figure 2B), indicating the absence of non-specific (e.g., motoric) effects.

#### Spontaneous withdrawal: ICSS

Baseline ICSS thresholds did not differ between groups (Table S1). ICSS threshold data during hours 2 (i.e., acute effect of MOR itself) and hours 6 – 98 (i.e., withdrawal period) of spontaneous withdrawal did not differ between the two groups receiving MOR (MOR + NX and MOR + SAL groups) or between the two groups receiving SAL (SAL + SAL and SAL + NX groups). Therefore, data from these groups were combined into single MOR (n = 37) and SAL (n = 20) groups for further analysis. Welch’s corrected t-test showed no significant difference in ICSS thresholds between MOR and SAL rats 2 hours after injection (MOR: 107.7 ±5.35%; SAL: 100.4 ± 2.50%), indicating that MOR iself did not affect ICSS. Two-way ANOVA on ICSS thresholds during spontaneous withdrawal 6-98 hours after morphine injection revealed a significant main effect of time (F (7, 385) = 4.831, p < 0.0001) and group (morphine vs. saline) (F (1, 55) = 4.012, p = 0.05), but no significant interaction (Figure 2C). After correcting for multiple comparisons, ICSS did not significantly differ between groups at any individual time-point post injection. However, peak ICSS threshold values between hours 6 and 98 (regardless of the time point at which they occurred) differed significantly between the morphine (117.1 ± 2.15%) and saline (110.4 ± 1.63%) groups (Welch-corrected t(54.9) = 2.50, p = 0.02).

No significant differences were observed in ICSS response latencies between groups during baseline sessions (Table S1), 2 hours after injection (MOR: 103.2 ± 3.01%; SAL: 96.43 ± 1.82%), or during spontaneous withdrawal (Figure 2D).

#### Precipitated and spontaneous withdrawal: Somatic signs

Somatic withdrawal signs were significantly elevated in the MOR + NX group during precipitated withdrawal, and in the MOR condition during spontaneous withdrawal (Supplementary Material, Figure S1).

### MSA in the Mor + NX Group

#### Acquisition (n = 29)

Two-way ANOVA revealed a significant main effect of lever (active vs inactive) (F (1, 28) = 58.38, p < 0.0001) and session (F (9, 252) = 2.896, p = 0.003) on responses during the first 10 days of acquisition (Figure 3A). Sidak’s multiple comparison test showed significantly higher responses on the active lever during all acquisition sessions (all t ≥ 4.88, all p < 0.0001).

**Figure 3.**
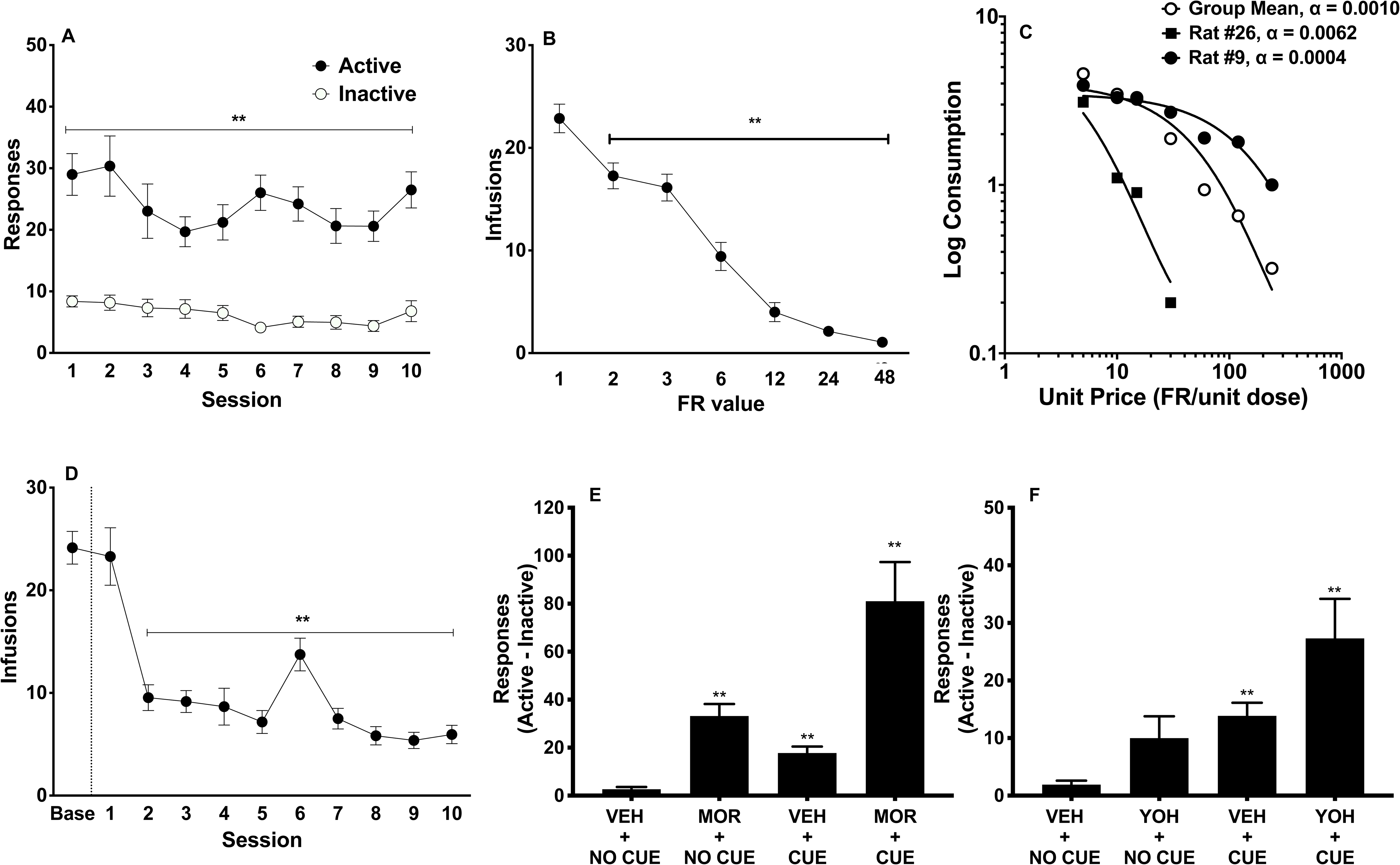
MSA in MOR + NX rats. (A) Mean (± SEM) active and inactive lever presses during the first 10 sessions of acquisition. ** Different compared to inactive lever presses, p < 0.01. (B) Mean (± SEM) infusions at each FR during demand testing. ** Different compared to infusions at FR1, p < 0.01. (C) Exponential demand curve describing morphine consumption as a function of unit price for rats as a group, and for individual rats with relatively high (rat #26) and low (rat #9) elasticity of demand (α). (D) Mean (± SEM) infusions during baseline and during the first 10 extinction sessions, ** Different compared to pre-extinction (baseline), p < 0.01. The increase in infusion rates during session 6 reflects spontaneous recovery following the weekend break in extinction testing. Mean (± SEM) reinstatement scores (differences between active and inactive lever responses) during morphine- and cue-induced reinstatement (E) and yohimbine-and cue-induced reinstatement (F). ** Different compared to VEH + NO CUE, p < 0.01.

#### Demand (n = 25)

Increases in FR requirement resulted in a progressive reduction in morphine consumption (Figure 3B). A one-way repeated measures ANOVA revealed a significant effect of FR on number of morphine infusions (F (3.59, 82.63) = 79.43, p < 0.0001). A post-hoc Dunnett’s multiple comparisons test showed that infusions at all subsequent FRs were significantly lower than at FR 1 (all q ≥ 4.23, all p ≤ 0.002). Morphine consumption during demand testing was well-described by an exponential demand function, with R^2^ values typically ≥ 0.80 for individual animals (Table S2) and R^2^ = 0.94 for rats as a group, with considerable individual variability in elasticity of demand (see Figure 3C).

#### Extinction (n = 24)

Repeated-measures ANOVA revealed a significant effect of session on infusion rates during extinction (F (3.64, 83.61) = 38.12, p > 0.001; Figure 3D). A post-hoc Dunnett’s test showed that infusions at all sessions after session 1 were significantly reduced compared to pre-extinction (baseline) levels (all p < 0.01).

#### Morphine- and cue-induced reinstatement (n = 23)

A one-way repeated measures ANOVA of reinstatement scores during morphine- and cue-induced reinstatement showed an overall effect of treatment (F(1.12, 24.73) = 18.18, p < 0.001) (Figure 3E). The Holm-Sidak’s multiple comparisons test revealed that the MOR + NO CUE, VEH + CUE and MOR + CUE conditions all resulted in significantly higher reinstatement scores compared to the VEH + NO CUE condition (all t ≥ 4.82, all p < 0.001). MOR+CUE resulted in significantly higher reinstatement than VEH + CUE (t = 3.83, p = 0.002) and MOR + NO CUE (t = 3.80, p = 0.002).

#### Stress- and cue-induced reinstatement (n = 22)

There was an overall effect of treatment on reinstatement scores during stress-and cue-induced reinstatement (F(1.68, 35.35) = 8.92, p = 0.001) (Figure 3F). The Holm-Sidak’s multiple comparisons test revealed that VEH + CUE (t = 5.98, p < 0.001) and YOH + CUE treatment (t = 3.75, p = 0.006) resulted in higher reinstatement scores than the VEH + NO CUE control condition, whereas the YOH + NO CUE condition did not. Responding during YOH + CUE reinstatement was significantly higher than during the YOH + NO CUE condition (t = 3.17, p = 0.02), but did not differ from the VEH + CUE condition.

### Correlations in the MOR + NX group

Composite z-scores for ICSS thresholds during precipitated withdrawal were significantly correlated with all individual measures: peak ICSS threshold (r = 0.92, p < 0.001), average ICSS threshold (r = 0.86, p < 0.001) and degree of sensitization of WIA (r = 0.54, p = 0.003). This validates our use of z-scores to measure cumulative precipitated withdrawal severity [see 39, 40]. Pearson’s r revealed that greater composite WIA severity during precipitated withdrawal correlated with lower infusions during acquisition of MSA and higher elasticity of demand (i.e., lower reinforcing efficacy) (Table 1, Figure S2). Greater peak ICSS threshold elevation during spontaneous withdrawal was associated with lower acquisition and reinstatement induced by morphine alone or morphine + cue (Table 1, Figure S3), and marginally correlated with elasticity of demand (p = 0.06, Figure S3). Most MSA measure correlated significantly with at least one other MSA measure (Table 1). However, no individual MSA measure correlated with as wide a range of other MSA measures as did WIA (Table 1).

**Table 1.**

| Variables | 1.<br>Precip<br>WIA | 2.<br>Spont.<br>WIA | 3.<br>Acquisition | 4.<br>Log $\alpha$ | 5.<br>Extinction | 6.<br>MOR+<br>CUE | 7.<br>MOR+<br>NO CUE | 8.<br>YOH+<br>CUE | 9.<br>YOH +<br>NO CUE |
| --- | --- | --- | --- | --- | --- | --- | --- | --- | --- |
| 1. Precip. WIA | — |  |  |  |  |  |  |  |  |
| 2. Spont. WIA | .41* | — |  |  |  |  |  |  |  |
| 3. Acquisition | -.39* | -.55** | — |  |  |  |  |  |  |
| 4. Log $\alpha$ | .43* | .40 | -.42* | — | | | | | |
| 5. Extinction | .40 | -.05 | .04 | -.28 | — |  |  |  |  |
| 6. MOR + CUE | -.07 | -.52** | .25 | -.10 | .01 | — |  |  |  |
| 7. MOR + NO CUE | -.22 | -.59** | .28 | -.13 | .01 | .81*** | — |  |  |
| 8. YOH + CUE | -.32 | -.14 | .17 | -.55** | .31 | -.19 | -.15 | — |  |
| 9. YOH + NO CUE | .12 | .09 | -.14 | -.20 | .19 | -.23 | -.35 | .59** | — |
| 10. VEH + CUE | -.18 | -.02 | .09 | -.58** | .21 | -.19 | -.24 | .74*** | .58** |

Correlation between ICSS withdrawal measures and MSA measures (Pearson's R) in the Mor + NX group. 1. Standardized composite score of average WIA, peak WIA, and degree of sensitization of WIA during naloxone-precipitated withdrawal. 2. Peak ICSS threshold during spontaneous withdrawal. 3. Average infusions during first 10 days of acquisition. 4. Log-transformed elasticity of demand computed from exponential demand model. Higher log $\alpha$ = lower reinforcement efficacy. 5. Average infusions during first 10 days of extinction; 6.-10. Reinstatement score (active – inactive lever pressing) during reinstatement induced by morphine with cue (MOR+CUE), morphine with no cue (MOR+ NO CUE), yohimbine with cue (YOH+CUE), yohimbine with no cue (YOH+ NO CUE) and cue alone (averaged across VEH + CUE condition during morphine- and yohimbine-induced reinstatement testing). \*\*\*, \*\*\*, p ≤ 0.05, 0.01, 0.001.

### Secondary correlations in the MOR + NX group

There were no significant correlations between any of the secondary predictors (i.e., somatic signs during acute dependence testing, locomotor activity) and any measure of MSA (all p-values > 0.05, data not shown).

### MSA in control groups

Rats in the control groups did not differ from the MOR + NX group on any primary MSA measure (see Supplementary Materials, Figure S4).

### Late-stage dependence

ICSS thresholds were significantly elevated during precipitated withdrawal during late-stage dependence (Figure S5), while somatic signs were significantly elevated during spontaneous withdrawal (Figure S5). However, these effects were not correlated with most MSA measures (see Supplementary Materials)

## 4. Discussion

Greater WIA during antagonist-precipitated and spontaneous withdrawal in an acute dependence model was associated with lower vulnerability on multiple measures of subsequent i.v. MSA (e.g., elasticity of demand, reinstatement). In fact, WIA predicted a wider range of MSA measures than did any individual measure of MSA. These findings are consistent with the principle that initial drug sensitivity is an important predictor of subsequent drug use [e.g., 1, 2, 3], and also support the notion that drug withdrawal sensitivity may be protective against drug addiction [17, 19, 47]. Our findings with outbred rats also complement findings of lower WIA in rats bred for high saccharin consumption [19], a line that exhibits greater SA of opioids and other drugs [23]. Together, these data identify WIA as a potential target for understanding behavioral and neurobiological mechanisms underlying the emergence of opioid addiction.

Several features of WIA during acute dependence distinguish it from other behavioral measures of addiction vulnerability. First WIA is unique in that it reliably predicts individual differences in opioid SA in outbred rats, whereas numerous established behavioral markers of individual differences in stimulant and alcohol SA (e.g., sensation-seeking as measured by open-field locomotor activity, impulsivity) do not [35, 48, 49]. Indeed, open-field activity was not correlated with any measure of MSA in this study, consistent with our previous findings using a more limited set of MSA measures [35]. WIA also differs from other behavioral predictors of SA of other drugs in that it predicted a variety of measures of drug SA rather than just a limited few [e.g., 4, 5]. An additional unique feature of WIA is that it is an outcome of early opioid exposure, as opposed to a preexisting disposition. Therefore, WIA may represent a neuroadaptive *mechanism* underlying addiction vulnerability, as opposed to only a behavioral *indicator*. As such, WIA promises to provide unique information on addiction vulnerability to complement findings obtained using existing behavioral markers of addiction vulnerability.

In contrast to WIA, somatic signs during acute dependence did not predict any primary MSA measure. These data complement previous findings indicating that affective/emotional and somatic withdrawal signs are mediated by distinct neurobiological mechanisms [50, 51], and supports the notion that the former have greater relevance to addiction vulnerability [52].

The current findings contrast with some studies reporting a *positive* relationship between withdrawal sensitivity and addiction vulnerability [52–55]. Numerous methodological differences between studies could account for this discrepancy (e.g., drug class studied, etc). In addition, the current acute dependence model isolates the earliest stages of dependence, while prior studies involved subjects in which dependence had already been established. As such, withdrawal sensitivity may shift from being a protective factor to a vulnerability factor for addiction as dependence develops [56]. The fact that WIA during late stage dependence was associated with *greater* resistance to extinction (see Supplemental Materials) may be consistent with this possibility. The lack of correlation between WIA during acute dependence and late-stage dependence (Supplementary Material) also suggests that each of these stages may provide unique information. Use of larger group sizes in order to provide adequate statistical power is needed to further address this issue, which was not a primary goal of this study.

Comparison of MSA in the MOR + NX and control groups suggests that a prior history of morphine exposure and/or withdrawal had limited effects on subsequent MSA. These data contrast with findings that repeated, experimenter-administered acute drug injections (and presumably spontaneous withdrawal episodes) enhance subsequent drug SA [33, 57, 58]. However, this phenomenon has been reported using stimulants rather than opioids and has involved a larger number of experimenter-administered injections than were used here. These or other methodological differences across studies (e.g., duration of interval between the final acute injection and onset of SA) may account for the similar MSA across groups in this study.

Behavioral economics has been useful for understanding individual differences in addiction vulnerability in both humans and animals, but has not been applied extensively to MSA. Consistent with our previous study [35], an exponential demand function generally provided a good fit for morphine consumption under an FR escalation procedure. There were also considerable individual differences in α (reinforcing efficacy) that were correlated with severity of WIA during precipitated withdrawal, and a similar trend was observed for spontaneous withdrawal. Together, these data further support the utility and sensitivity of behavioral economics to study individual differences in opioid addiction vulnerability.

In conclusion, this study establishes WIA as one of the first behavioral measures to reliably predict individual differences in future opioid SA. Future studies characterizing the neurobiological mechanisms underlying WIA will allow for identification of addiction-related effects (i.e., those uniquely related to severity of WIA) from other, corollary effects of opioids. Hence, further use of this model in rats promises to provide fresh insights into neurobiological mechanisms underlying vulnerability and/or resilience to opioid addiction.

## Acknowledgements

The authors thank Mary Krueger, Joseph Tombers, Haley Rudnick, and Nettie Enshayan for their excellent technical assistance.

## Supplementary Material

### Animals

Male adult Sprague Dawley rats (Harlan/Envigo, Indianapolis, IN) weighing 276-300 g at arrival were used. All rats were individually housed in a temperature- and humidity-controlled colony room with unlimited access to water under a reversed 12-h light/dark cycle (lights off at 10:00 hr). All behavioral testing occurred during the dark (active) phase. Beginning one week following arrival, food was restricted to 18 g/day to facilitate operant performance, avoid detrimental health effects of long-term ad libitum feeding, and limit catheter migration.

### Apparatus

#### Intracranial self-stimulation (ICSS)

Rats were tested in operant conditioning chambers (29×26×33 cm; Med Associates, St. Albans, VT, USA) placed inside sound-attenuating cubicles. A 5-cm-wide metal wheel manipulandum was fixed to the front wall. Brain stimulation was administered with constant current stimulators (model #PHM-152, Med Associates). Rats were connected to the stimulation circuit through bipolar leads (Plastics One, Roanoke, VA, USA) attached to gold-contact swivel commutators (Plastics One). MED-PC IV software was used to control stimulation parameters and for data collection.

#### Morphine self-administration (MSA)

MSA sessions were conducted using 16 standard operant conditioning chambers (model ENV-007, Med Associates, Inc). Each chamber contained two response levers, a white stimulus (i.e., cue) light located 2 cm above each lever, and a house light that provided ambient illumination. Each chamber was placed inside a sound-attenuating cubicle equipped with an exhaust fan that provided masking noise. An infusion pump (model PHM-100-15, Med Associates) placed outside each cubicle delivered infusions in a volume of 0.1 ml/kg over approximately 1 second. MED-PC IV software (Med Associates) was used for operating the experimental apparatus and recording data.

### Surgery

#### ICSS

Animals were anesthetized with ketamine (75 mg/kg, i.m.) and dexmedetomidine (0.5 mg, i.m.) and implanted with a bipolar stainless steel electrode (Plastics One) in the medial forebrain bundle at the level of the lateral hypothalamus as described in [1]. Animals were allowed to recover for at least 1 week prior to ICSS training. During the first 2 days of recovery, all animals received injections of the antibiotic ceftriaxone (5.25 mg, i.m.) and the analgesic buprenorphine (0.1 mg/kg, s.c.).

#### MSA

Each rat was implanted with a chronic indwelling catheter into the right jugular vein under isoflurane (1-3%) anesthesia, using general surgical procedures described in detail elsewhere [2, 3]. The catheter was externalized between the scapulae and attached to a vascular-access harness (VAH95AB, Instech Laboratories, Plymouth Meeting, PA) that allowed connection to a fluid swivel via a tether for morphine administration. Animals were allowed to recover for one week after surgery, during which time they received daily i.v. infusions of heparinized saline, ceftriaxone antibiotic (5.25 mg, first three days only), and s.c. injections of buprenorphine (0.05 mg/kg; first two days only) for analgesia. Infusions of methohexital (0.1 ml, 10 mg/ml, i.v.) were administered to check catheteer patency post-session on Fridays. If a catheter became occluded (indicated by a failure of the animal to exhibit anesthesia within 3-5 sec after methohexital infusion), another catheter was implanted into the ipsilateral femoral vein. Failure of this second catheter resulted in removal of the animal from the study.

### General testing procedures

#### ICSS

Each trial was initiated with presentation of a non-contingent stimulus (0.1-ms cathodal square wave pulses at a frequency of 100 Hz for 500 ms) followed by a 7.5-s window, during which a positive response on the wheel manipulandum produced a second contingent stimulation identical to the first. Lack of responding during the 7.5-s window was considered a negative response. Each positive or negative response was followed by a variable inter-trial interval averaging 10 s (range, 7.5–12.5 s), during which time additional responses delayed the onset of the subsequent trial by 12.5 s. Stimulus intensities were presented in four alternating descending and ascending series (step size, 5 μA), with five trials presented at each current intensity step. The current threshold for each series was defined as the midpoint between two consecutive intensity steps that yielded three or more positive responses and two consecutive intensity steps that yielded three or more negative responses. The overall ICSS threshold for the session was defined as the mean of the current thresholds from the four alternating series. To assess performance effects (e.g., motor disruption), response latencies (time between onset of the non-contingent stimulus and a positive response) were averaged across all trials in which a positive response was made.

#### MSA

During each 2 hr session, responding on the left (“active”) response lever resulted in an i.v. infusion of morphine sulfate (0.2 mg/kg/inf) that was accompanied by offset of the house light and the onset of a white cue light above the active response lever. Following a 30-second timeout period, the cue light above the active lever was extinguished to signal availability of the next infusion. Responses on the other response lever (the “inactive” lever) were recorded but had no programmed consequences. This unit dose and access duration support reliable MSA in the absence of self-mutilation associated with higher unit doses and longer sessions [32]. On the first day of MSA (always a Friday), food powder was placed on the active lever to facilitate contact with the lever. Data from this session were not included in the data analysis.

#### Somatic withdrawal signs

Rats were placed in a clear plastic circular chamber and recorded with a digital camera for 10 min. Recordings were later scored for somatic signs by a blinded trained observer using a validated checklist [41]. Individual categories of withdrawal signs included eye blinks, wet dog shake, escape jumps, abdominal constrictions, swallowing movement, facial fasciculations, abnormal posture, ptosis, penile grooming, chromodacryorrhea, salivation and diarrhea.

## Supplementary Results

### Acute dependence: Somatic withdrawal signs

One-way ANOVA on total somatic signs during the 5^th^ session of naloxone-precipitated withdrawal indicated a significant effect of group (F (3, 52) = 11.51, p < 0.0001). Dunnett’s multiple comparison test revealed significantly higher scores in the the MOR + NX group compared to the SAL + SAL group (q(52) =2.62, adjusted p = 0.03). In contrast, neither the MOR + SAL or SAL + NX group differed significantly from the SAL + SAL group (Figure S1A).

During spontaneous withdrawal, somatic sign scores did not differ between the two groups receiving MOR (MOR + NX and MOR + SAL groups) or between the two groups receiving SAL (SAL + SAL and SAL + NX groups). Data from these groups were therefore combined into single MOR (n = 37) and SAL (n = 20) conditions. T-tests showed that scores were significantly higher in the MOR condition compared to the SAL condition (Welch-corrected t(51.31) = 2.25, p = 0.03) (Figure S1B).

### Correlations in the MOR + NX group

Figures S2 and S3 show scatterplots portraying the relationship between composite precipitated WIA (Figure S2) and peak spontaneous WIA (Figure S3) during acute dependence testing and various MSA measures.

### Comparison of MSA in the MOR + NX and control groups

Rats in the MOR + NX group and control groups (MOR + SAL, SAL + NX, SAL + SAL) did not differ on most measures of MSA. Two-way repeated measures ANOVA on infusion rates between the MOR + NX and the control groups during the first 10 days of acquisition indicated a significant overall effect of session (F (9, 468) = 2.20, p = 0.02) but no effect of group or interaction between group and session (Figure S4A). Two-way ANOVA on group and FR during demand testing showed a significant main effect of FR (F (6, 246) = 152, p < 0.001) and interaction between group and FR (F (18, 246) = 2.144, p = 0.005). but no main effect of group (Figure S4B). The MOR + NX had significantly lower infusions at FR 2 and FR 3 compared to the SAL + SAL group (all p < 0.01). However, a one-way ANOVA comparing elasticity of demand (α) (i.e., the primary outcome) indicated no significant difference between groups (Figure S4C). The MOR + NX and control groups also did not significantly differ in three secondary behavioral economic measures: intensity of demand (Q_0_), maximal response output (O_max_) and the unit price at which maximal response output occurred (P_max_) (Table S2). Additionally, the groups did not differ in infusions during extinction, as a two-way repeated measures ANOVA revealed a significant overall effect of session (F (10, 410) = 79.53, p < 0.0001) (Figure S4D) but no effect of group or group x session interaction. Finally, two-way ANOVAs showed a main effect of reinstatement condition on reinstatement scores during morphine- and cue-induced reinstatement (F (1.291, 51.65) = 28.95, P < 0.0001) and yohimbine- and cue-induced reinstatement (F (1.747, 65.79) = 22.68, P < 0.0001), with responses significantly elevated during all drug (morphine or yohimbine) and/or cue conditions compared to the VEH + CUE control condition (Figure S4E-F). However, there was no significant main effect of group or interaction during either morphine- and cue-induced reinstatement testing or yohimbine- and cue-reinstatement testing.

### Late-stage dependence

#### Precipitated withdrawal: ICSS

Baseline ICSS thresholds and response latencies for late-stage precipitated withdrawal did not differ between groups (Table S3). Two-way ANOVA on ICSS thresholds during precipitated withdrawal revealed a significant main effect of group (F (3, 22) = 6.2, p = 0.003), but no significant effect of session or interaction. Holm-Sidak’s multiple comparison showed significantly higher ICSS thresholds in the MOR + NX group compared to the SAL + SAL control group during all sessions (all t(22) ≥ 3.71, all p < 0.05) (Figure S5A). There was no significant effect of group, session, or group x session interaction on ICSS latencies during precipitated withdrawal (data not shown).

#### Spontaneous withdrawal: ICSS

Baseline ICSS thresholds and response latencies did not differ between groups (Table S3). ICSS thresholds during hours 2 (agonist effect) and hours 6 – 98 (withdrawal period) of spontaneous withdrawal did not differ between the two groups receiving MOR (MOR + NX versus MOR + SAL groups) or SAL (SAL + SAL versus SAL + NX groups). Data from these groups were therefore combined into a single MOR (n = 15) and SAL (n = 5) group. Welch’s corrected t test showed no significant difference in ICSS thresholds between MOR (118.4 ± 14.03%) and SAL (102.6 ± 2.58%) groups during the 2-hour session. Two-way ANOVA on ICSS thresholds during spontaneous withdrawal 6-98 hours after morphine injection revealed no significant main effect of time, group (morphine vs. saline) or interaction (Figure S5B). However, comparison of peak ICSS threshold values between 6 hours and 98 hours (regardless of the time point at which they occurred) differed significantly between the morphine (121.7 ± 4.78%) and saline (106.8 ± 2.04%) groups (Welch-corrected t(17.54) = 2.85, p = 0.01). No significant difference in ICSS response latencies was observed between groups 2 hours after injection (agonist effect) or 6-98 hours after injection (withdrawal effect) (data not shown).

#### Precipitated and spontaneous withdrawal: Somatic signs

One-way ANOVA revealed a significant overall difference in somatic signs scores between groups during the 5^th^ session of precipitated withdrawal during late stage dependence (F(3, 23) = 4.57, p = 0.01). However, Dunnett’s multiple comparison indicated that none of the groups differed significantly from the SAL + SAL group (Figure S5C).

During spontaneous withdrawal, somatic signs did not differ between the two groups receiving MOR or between the two groups receiving SAL. Data from these groups were therefore combined into single MOR (n = 15) and SAL (n = 5) conditions, respectively. Total somatic signs in the MOR condition were significantly higher than in the SAL condition (Welch-corrected t(21.44) = 2.19, p = 0.04) (Figure S5D).

### Correlations during late-stage dependence

In MOR + NX rats, ICSS thresholds or somatic signs scores during late-stage dependence during precipitated or spontaneous withdrawal did not correlate with any MSA measure, except for a significant correlation between higher composite z-score for ICSS thresholds during precipitated withdrawal and higher infusions during extinction (i.e., *greater* resistance to extinction; r = 0.61, p = 0.04).

Comparison of withdrawal severity during acute dependence and late-stage dependence in rats that completed both phases suggested that these measures were largely independent. Thus, ICSS thresholds during precipitated or spontaneous withdrawal during late-stage dependence did not correlate with these same ICSS measures during acute dependence (all r: −0.16 ≤ r ≤ −0.04, all p ≥ 0.57). Somatic signs during spontaneous withdrawal correlated significantly between late-stage dependence and acute dependence (r = 0.73, p = 0.007), but this correlation was not observed during precipitated withdrawal (r = 0.13, p = 0.45).

---

**Table S1.** Mean (±SEM) ICSS thresholds (in µA) and response latencies (in sec) in experimental groups during baseline sessions during acute dependence testing.

|  | Naloxone-precipitated Withdrawal |  | Spontaneous Withdrawal |  |
| --- | --- | --- | --- | --- |
| | Threshold (μA) | Latency (sec) | Threshold (μA) | Latency (sec) |
| MOR+NX | 131.5 ± 13.37 | 2.62 ± 0.10 | 132.5 ± 13.99 | 2.62 ± 0.09 |
| MOR+SAL | 123.3 ± 12.45 | 2.69 ± 0.20 | 127.0 ± 12.80 | 2.59 ± 0.13 |
| SAL+NX | 128.8 ± 15.50 | 2.50 ± 0.17 | 133.3 ± 17.90 | 2.60 ± 0.20 |
| SAL+SAL | 142.4 ± 25.14 | 2.65 ± 0.11 | 141.1 ± 25.31 | 2.71 ± 0.14 |

**Table S2.**
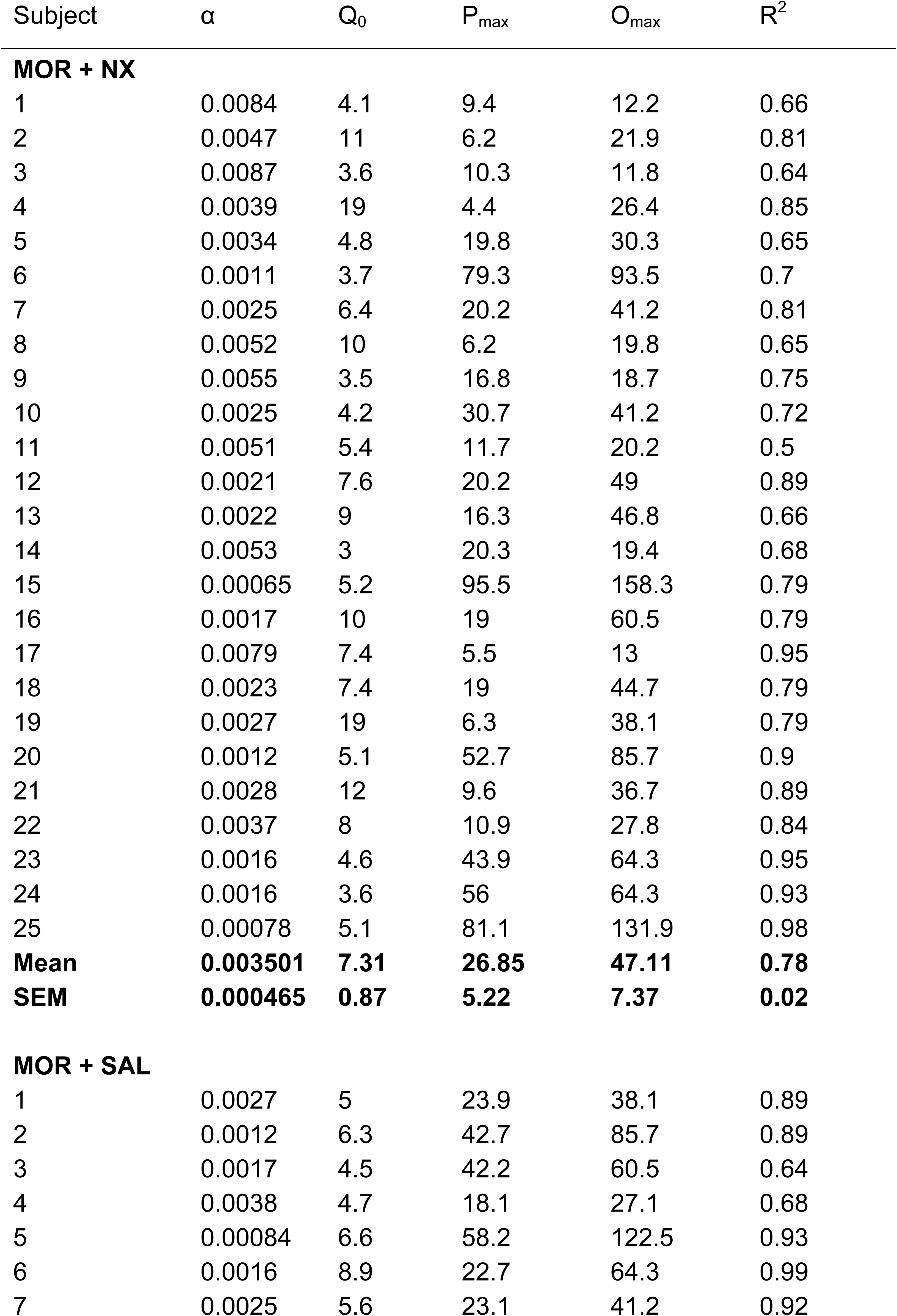
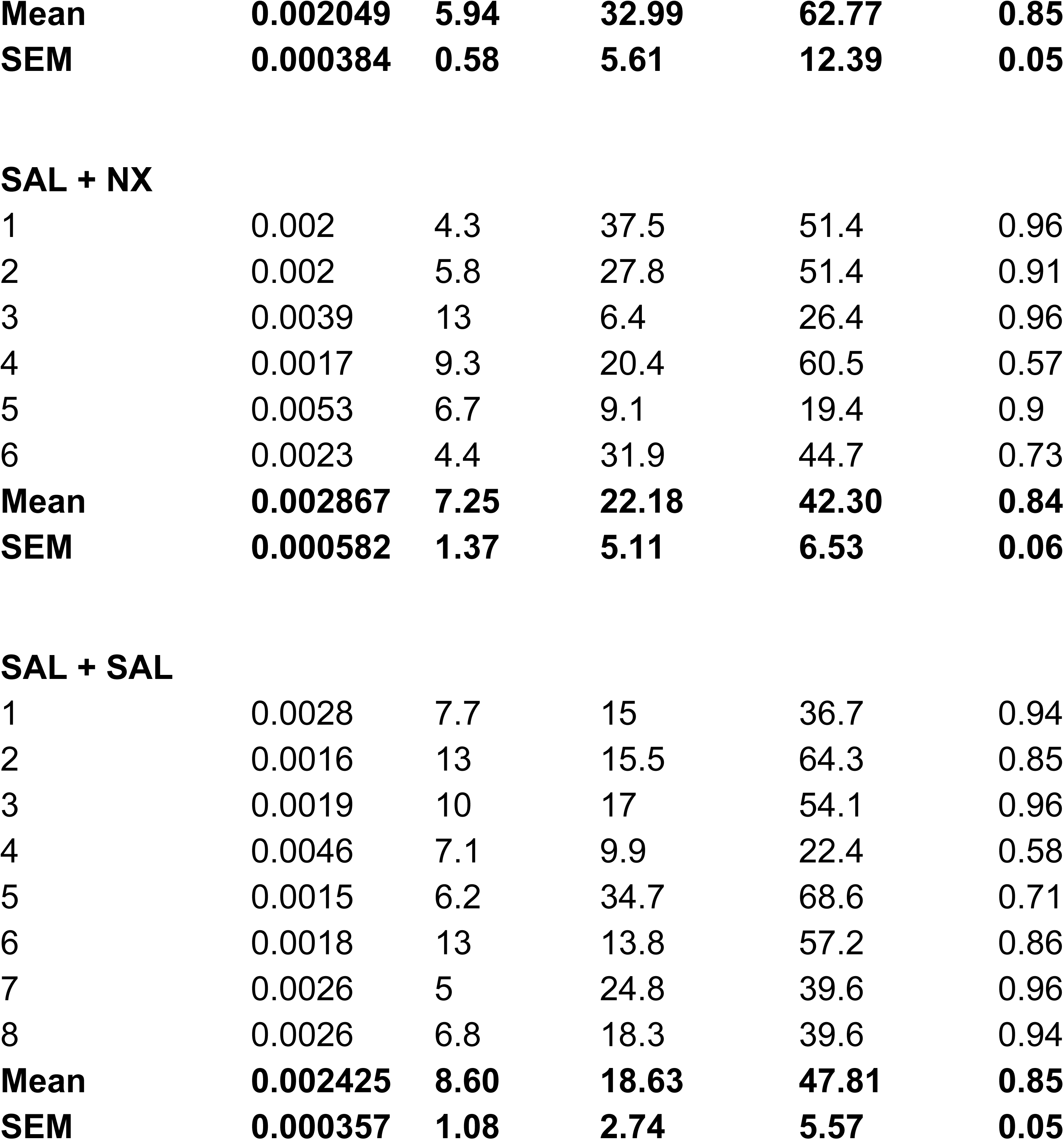
Exponential demand curve parameters for individual subjects. Note: the parameter k (range of consumption) is set to 1.8 log units.

| Subject | $\alpha$ | $Q_0$ | $P_{\max}$ | $O_{\max}$ | $R^2$ |
| --- | --- | --- | --- | --- | --- |
| **MOR + NX** |  |  |  |  |  |
| 1 | 0.0084 | 4.1 | 9.4 | 12.2 | 0.66 |
| 2 | 0.0047 | 11 | 6.2 | 21.9 | 0.81 |
| 3 | 0.0087 | 3.6 | 10.3 | 11.8 | 0.64 |
| 4 | 0.0039 | 19 | 4.4 | 26.4 | 0.85 |
| 5 | 0.0034 | 4.8 | 19.8 | 30.3 | 0.65 |
| 6 | 0.0011 | 3.7 | 79.3 | 93.5 | 0.7 |
| 7 | 0.0025 | 6.4 | 20.2 | 41.2 | 0.81 |
| 8 | 0.0052 | 10 | 6.2 | 19.8 | 0.65 |
| 9 | 0.0055 | 3.5 | 16.8 | 18.7 | 0.75 |
| 10 | 0.0025 | 4.2 | 30.7 | 41.2 | 0.72 |
| 11 | 0.0051 | 5.4 | 11.7 | 20.2 | 0.5 |
| 12 | 0.0021 | 7.6 | 20.2 | 49 | 0.89 |
| 13 | 0.0022 | 9 | 16.3 | 46.8 | 0.66 |
| 14 | 0.0053 | 3 | 20.3 | 19.4 | 0.68 |
| 15 | 0.00065 | 5.2 | 95.5 | 158.3 | 0.79 |
| 16 | 0.0017 | 10 | 19 | 60.5 | 0.79 |
| 17 | 0.0079 | 7.4 | 5.5 | 13 | 0.95 |
| 18 | 0.0023 | 7.4 | 19 | 44.7 | 0.79 |
| 19 | 0.0027 | 19 | 6.3 | 38.1 | 0.79 |
| 20 | 0.0012 | 5.1 | 52.7 | 85.7 | 0.9 |
| 21 | 0.0028 | 12 | 9.6 | 36.7 | 0.89 |
| 22 | 0.0037 | 8 | 10.9 | 27.8 | 0.84 |
| 23 | 0.0016 | 4.6 | 43.9 | 64.3 | 0.95 |
| 24 | 0.0016 | 3.6 | 56 | 64.3 | 0.93 |
| 25 | 0.00078 | 5.1 | 81.1 | 131.9 | 0.98 |
| **Mean** | **0.003501** | **7.31** | **26.85** | **47.11** | **0.78** |
| **SEM** | **0.000465** | **0.87** | **5.22** | **7.37** | **0.02** |
| **MOR + SAL** |  |  |  |  |  |
| 1 | 0.0027 | 5 | 23.9 | 38.1 | 0.89 |
| 2 | 0.0012 | 6.3 | 42.7 | 85.7 | 0.89 |
| 3 | 0.0017 | 4.5 | 42.2 | 60.5 | 0.64 |
| 4 | 0.0038 | 4.7 | 18.1 | 27.1 | 0.68 |
| 5 | 0.00084 | 6.6 | 58.2 | 122.5 | 0.93 |
| 6 | 0.0016 | 8.9 | 22.7 | 64.3 | 0.99 |
| 7 | 0.0025 | 5.6 | 23.1 | 41.2 | 0.92 |
| **Mean** | **0.002049** | **5.94** | **32.99** | **62.77** | **0.85** |
| **SEM** | **0.000384** | **0.58** | **5.61** | **12.39** | **0.05** |
| **SAL + NX** |  |  |  |  |  |
| 1 | 0.002 | 4.3 | 37.5 | 51.4 | 0.96 |
| 2 | 0.002 | 5.8 | 27.8 | 51.4 | 0.91 |
| 3 | 0.0039 | 13 | 6.4 | 26.4 | 0.96 |
| 4 | 0.0017 | 9.3 | 20.4 | 60.5 | 0.57 |
| 5 | 0.0053 | 6.7 | 9.1 | 19.4 | 0.9 |
| 6 | 0.0023 | 4.4 | 31.9 | 44.7 | 0.73 |
| **Mean** | **0.002867** | **7.25** | **22.18** | **42.30** | **0.84** |
| **SEM** | **0.000582** | **1.37** | **5.11** | **6.53** | **0.06** |
| **SAL + SAL** |  |  |  |  |  |
| 1 | 0.0028 | 7.7 | 15 | 36.7 | 0.94 |
| 2 | 0.0016 | 13 | 15.5 | 64.3 | 0.85 |
| 3 | 0.0019 | 10 | 17 | 54.1 | 0.96 |
| 4 | 0.0046 | 7.1 | 9.9 | 22.4 | 0.58 |
| 5 | 0.0015 | 6.2 | 34.7 | 68.6 | 0.71 |
| 6 | 0.0018 | 13 | 13.8 | 57.2 | 0.86 |
| 7 | 0.0026 | 5 | 24.8 | 39.6 | 0.96 |
| 8 | 0.0026 | 6.8 | 18.3 | 39.6 | 0.94 |
| **Mean** | **0.002425** | **8.60** | **18.63** | **47.81** | **0.85** |
| **SEM** | **0.000357** | **1.08** | **2.74** | **5.57** | **0.05** |

**Table S3.**
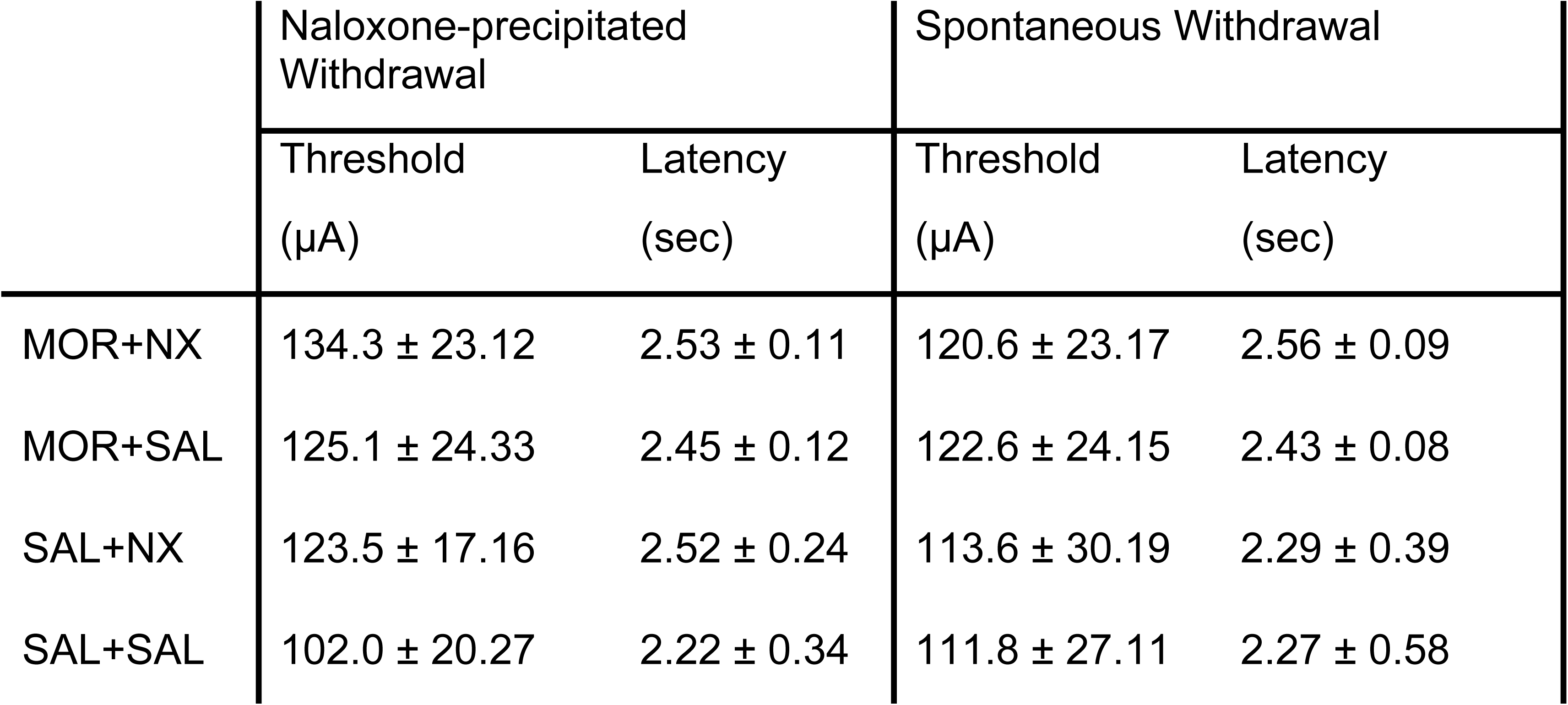
Mean (±SEM) ICSS thresholds (in µA) and response latencies (in sec) in experimental groups during baseline sessions during late-stage dependence withdrawal testing.

|  | Naloxone-precipitated Withdrawal |  | Spontaneous Withdrawal |  |
| --- | --- | --- | --- | --- |
| | Threshold (μA) | Latency (sec) | Threshold (μA) | Latency (sec) |
| MOR+NX | 134.3 ± 23.12 | 2.53 ± 0.11 | 120.6 ± 23.17 | 2.56 ± 0.09 |
| MOR+SAL | 125.1 ± 24.33 | 2.45 ± 0.12 | 122.6 ± 24.15 | 2.43 ± 0.08 |
| SAL+NX | 123.5 ± 17.16 | 2.52 ± 0.24 | 113.6 ± 30.19 | 2.29 ± 0.39 |
| SAL+SAL | 102.0 ± 20.27 | 2.22 ± 0.34 | 111.8 ± 27.11 | 2.27 ± 0.58 |

**Figure S1:**
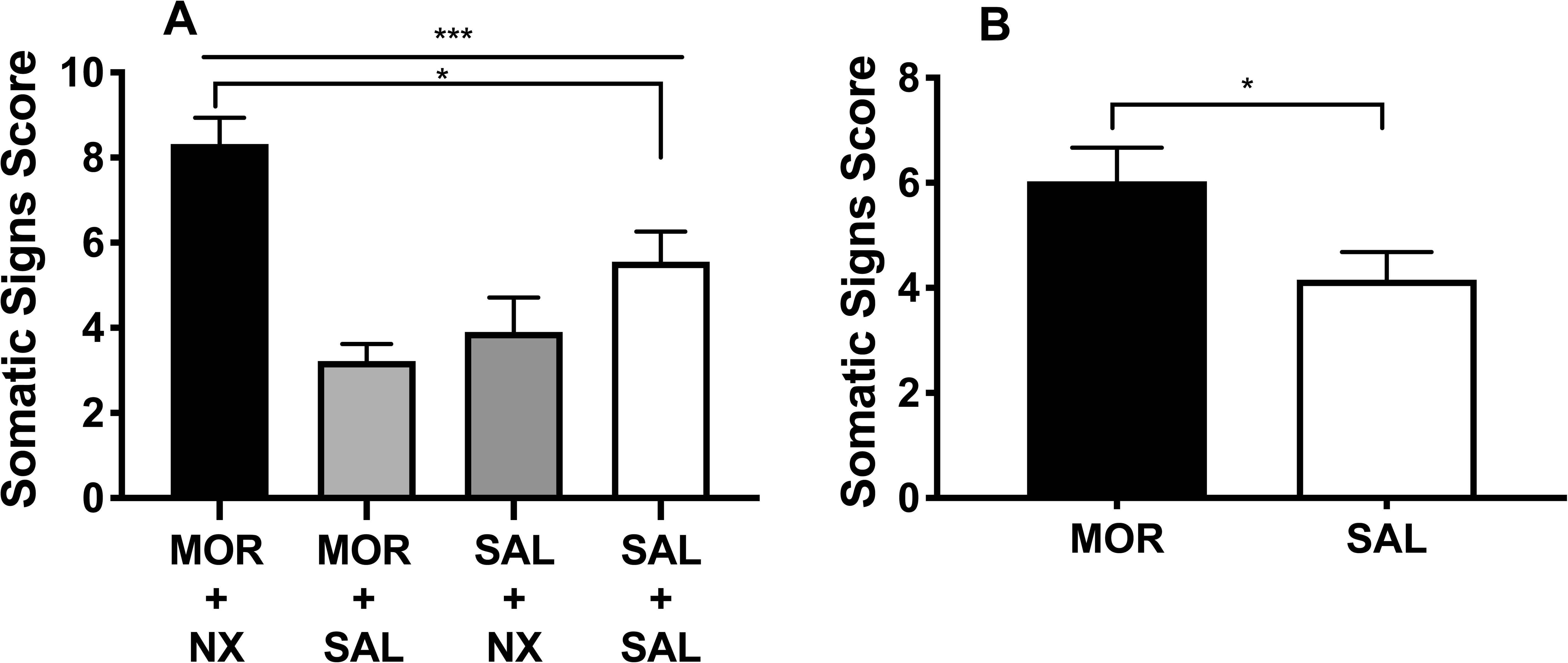
(A) Mean (± SEM) somatic signs in groups on the 5^th^ day of acute dependence precipitated withdrawal testing. (B) Mean (± SEM) somatic signs in the MOR and SAL condition 26 hours after injection during acute dependence spontaneous withdrawal testing. *** Significant effect of group, p < 0.001. * Different from SAL + SAL group or SAL condition, p < 0.05.

**Figure S2:**
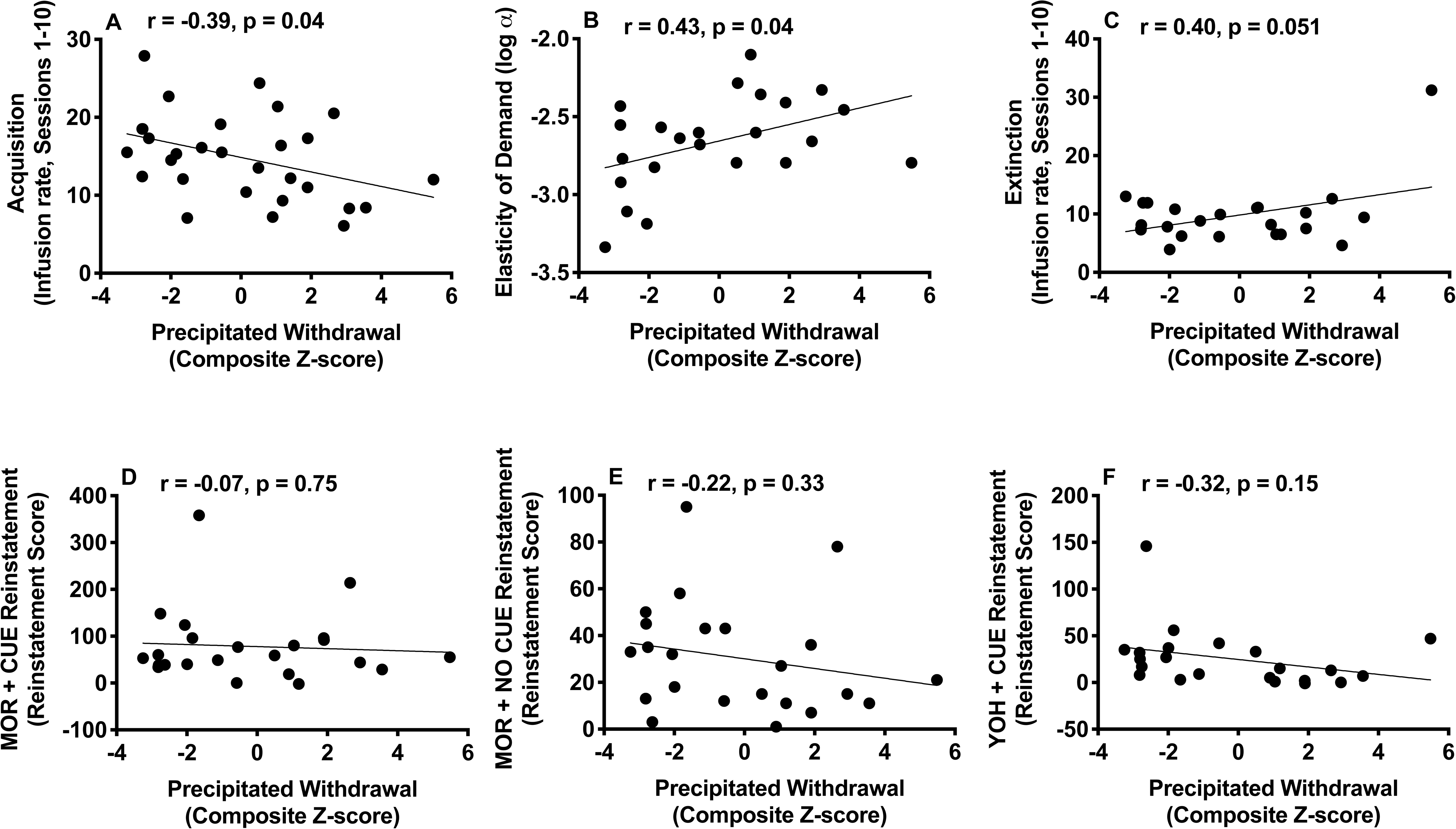
Scatterplots with regression line depicting the bivariate relationships between ICSS precipitated withdrawal composite z-score and infusions during first 10 sessions of acquisition (A), log α (B), infusions during first 10 sessions of extinction (C), and reinstatement score during MOR + CUE (D), MOR + NO CUE (E) and YOH + CUE (F) reinstatement. Higher log α (elasticity of demand) = lower reinforcement efficacy.

**Figure S3:**
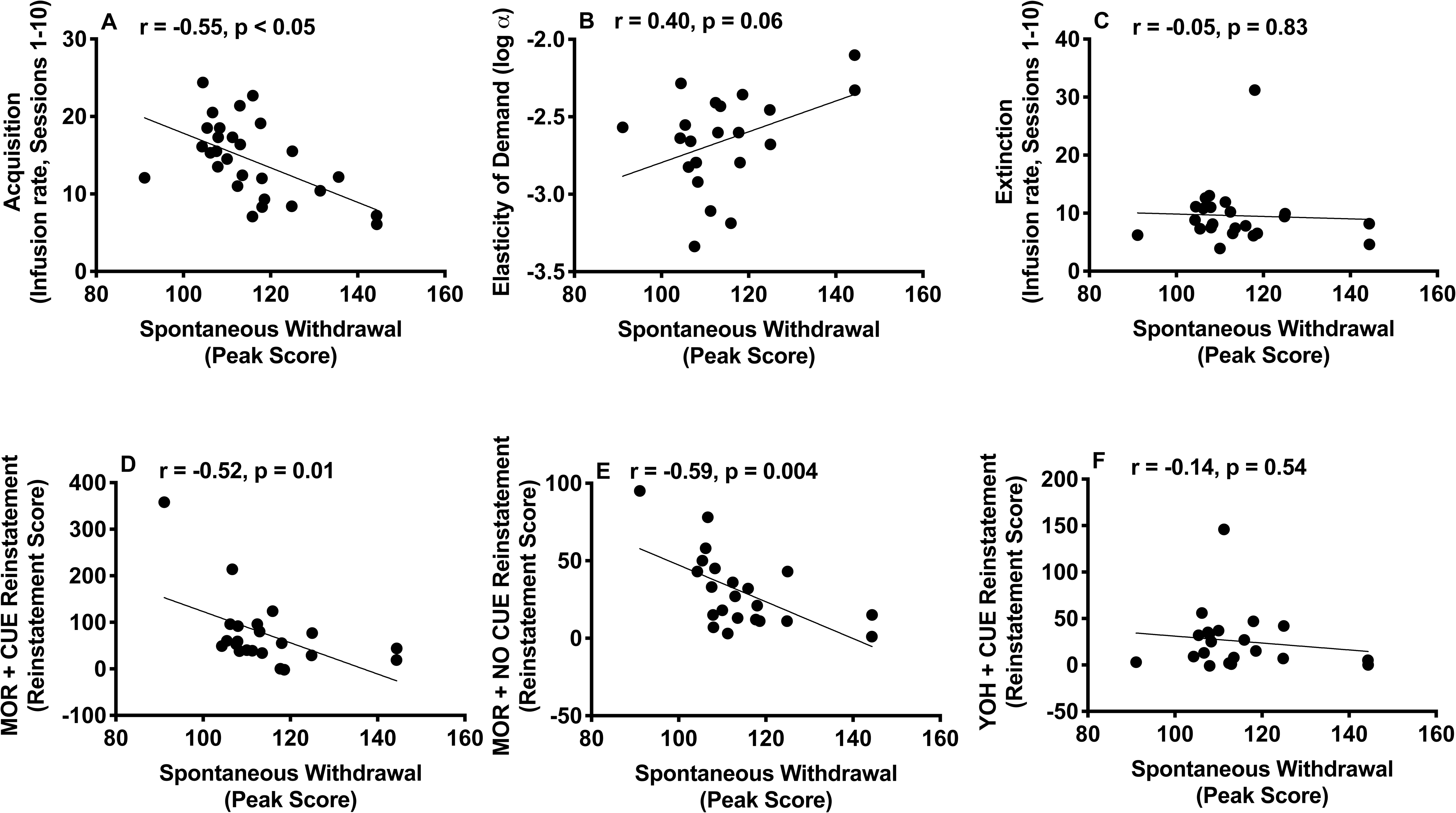
Scatterplots with regression line depicting the bivariate relationships between peak spontaneous withdrawal during ICSS testing and infusions during first 10 sessions of acquisition (A), log α (B), infusions during first 10 sessions of extinction (C), and reinstatement score during MOR + CUE (D), MOR + NO CUE (E) and YOH + CUE (F) reinstatement. Higher log α (elasticity of demand) = lower reinforcement efficacy.

**Figure S4:**
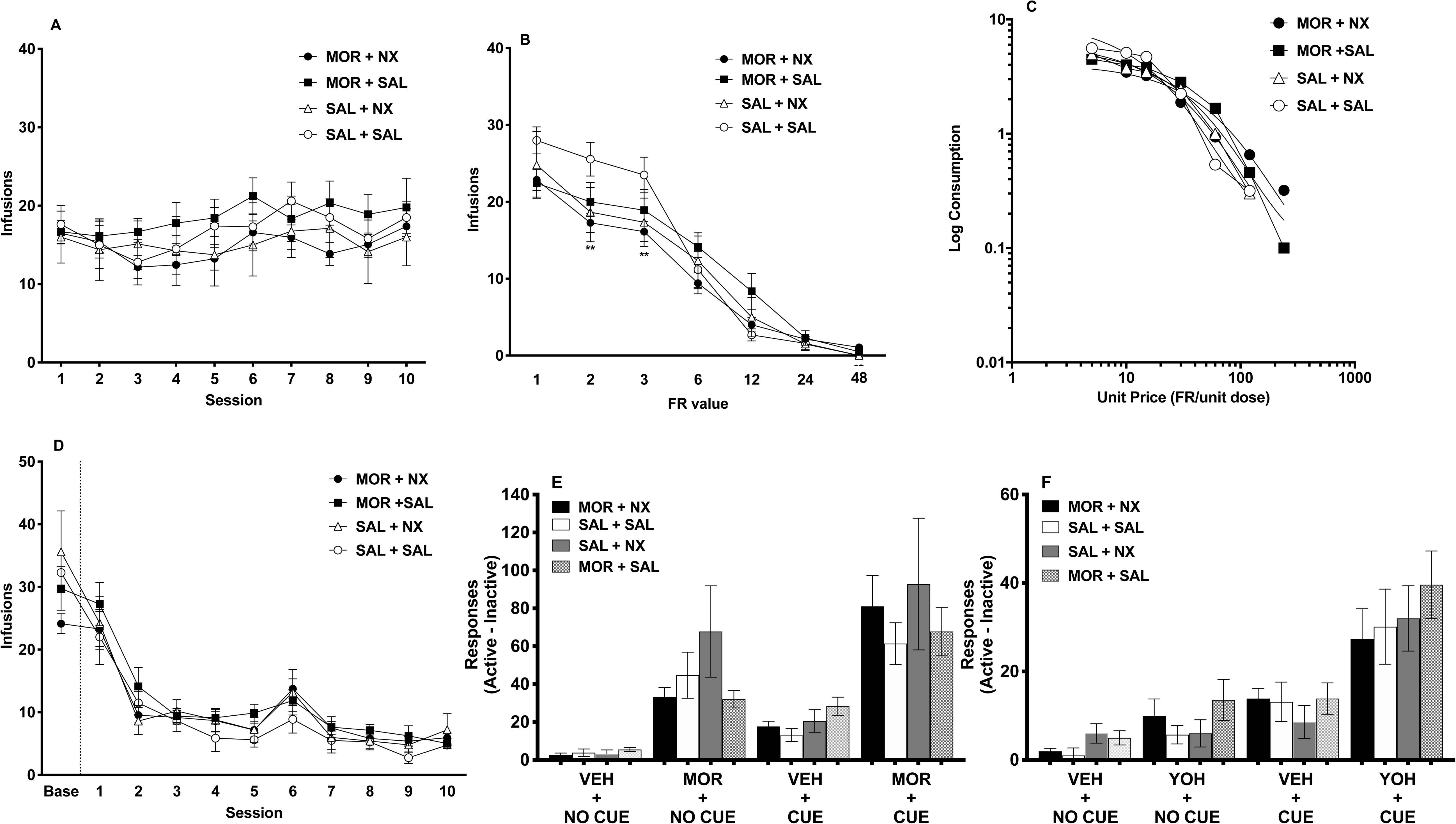
(A) Mean (± SEM) infusions during MSA acquisition for the MOR + NX and control groups. (B) Mean (± SEM) infusions at each FR during demand testing. ** Different from SAL + SAL group at that FR, p < 0.01. (C) Exponential demand curve describing morphine consumption as a function of unit price for rats as a group for each of the 4 groups. (D) Mean (± SEM) number of infusions during baseline and MSA extinction over 10 sessions. (E and F) Mean (± SEM) reinstatement scores during morphine- and cue-induced reinstatement (E) and yohimbine- and cue-induced reinstatement (F).

**Figure S5:**
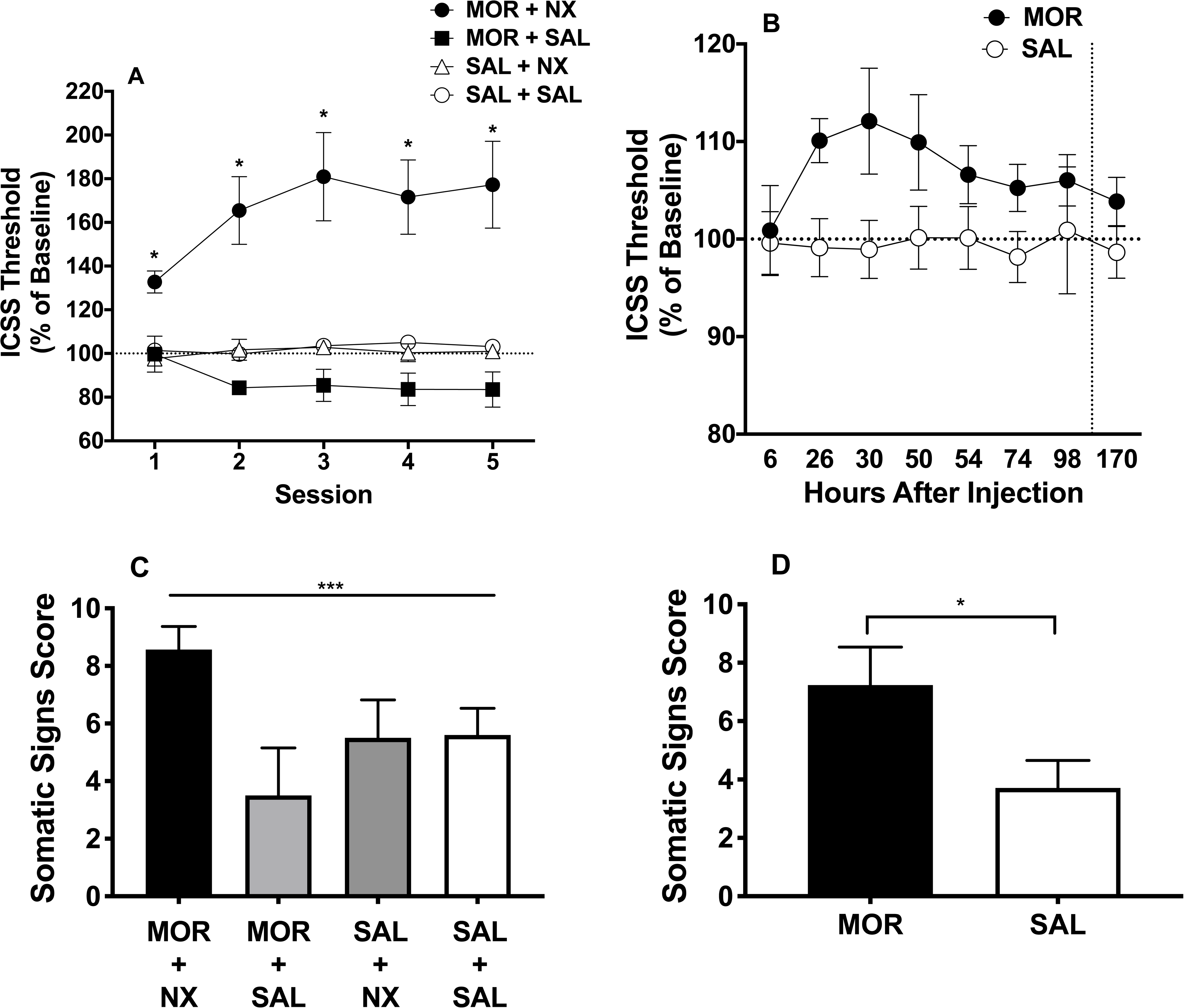
Mean (± SEM) ICSS thresholds (expressed as percent of baseline) during late-stage naloxone-precipitated withdrawal (A) and spontaneous withdrawal (B). * Different compared to SAL + SAL group at that session, p < 0.05. Mean (± SEM) somatic signs in groups on the 5th day of late-stage dependence precipitated withdrawal (C) and 26 hours after injection during late-stage dependence spontaneous withdrawal (D). *** Significant effect of group, p < 0.001. * Different from SAL condition, p < 0.05.

